# Species Composition and Distribution of Anopheles gambiae Complex Circulating in Kinshasa

**DOI:** 10.1101/2023.10.26.564181

**Authors:** Josue Zanga, Emery Metelo, Nono Mvuama, Victoire Nsabatien, Vanessa Mvudi, Degani Banzulu, Osée Mansiangi, Maxwel Bamba, Narcisse Basosila, Rodrigue Agossa, Roger Wumba

**Affiliations:** University of Kinshasa, Tropical Medicine Department, Kinshasa, Democratic Republic of the Congo.; Kinshasa School Public Health, Laboratory of Bio-ecology and Vector Control, Department of Health-Environment, Kinshasa, Democratic Republic of the Congo; University of Bandundu, Faculty of Medicine, Bandundu Ville, Democratic Republic of the Congo; University of Kinshasa, Department of Neurology, Kinshasa, Democratic Republic of the Congo; National Malaria Control Programme, Vector Control Service, Kinshasa, Democratic Republic of Congo; Cotonou Entomological Research Center (CREC), Cotonou, Benin

**Author notes:** **Corresponding authors**: Josue Zanga ( /+243815108117), Victoire Nsabatien (Victoire Nsabatien /+243829494537). **Mail list** Josue Zanga, Emery Metelo Nono Mvuama Victoire Nsabatien Vanessa Mvudi Degani Banzulu Osee Mansiangi, Maxwel Bamba Narcisse Basosila Fiacre Agossa, Roger Wumba.

**Keywords:** *Anopheles gambiae* complex, Composition, Distribution, Kinshasa, PCR

## Abstract

Understanding the distribution of *Anopheles* species in a region is an important task in the planning and implementation of malaria control programmes. This study was proposed to evaluate the composition and distribution of cryptic species of the main malaria vector, *Anopheles gambiae* complex, circulating in different districts of Kinshasa.

To study the distribution of members of the *An*. *gambiae* complex, *Anopheles* were sampled by CDC light trap and larva collection across the four districts of Kinshasa city between July 2021 and June 2022. After morphological identification, an equal proportion of *Anopheles gambiae* s.l. sampled per site were subjected to polymerase chain reaction (PCR) for identification of cryptic *An*. *gambiae* complex species.

The *Anopheles gambiae* complex was widely identified in all sites across the city of Kinshasa, with a significant difference in mean density, captured by CDC light, inside and outside households in Kinshasa (p=0.002). Two species of this complex circulate in Kinshasa: *Anopheles gambiae* (82.1%) and *Anopheles coluzzii* (17.9%). In all study sites, *Anopheles gambiae* was the most prevalent species. *Anopheles coluzzii* was very prevalent in Tshangu district. No hybrids (*Anopheles coluzzii*/*Anopheles gambiae*) were identified.

Two cryptic species of the *Anopheles gambiae* complex circulate in Kinshasa. *Anopheles gambiae* s.s., present in all districts and *Anopheles coluzzii*, with a limited distribution. Studies on the ecology of the larval sites are essential to better understand the factors influencing the distribution of members of the *An. gambiae* complex in this megalopolis.

## Context

The genus *Anopheles* is by far the most incriminated Culicidae vector targeted by vector control efforts. It is the vector responsible for the transmission of malaria, which is an infectious disease caused by protozoan parasites of the genus *Plasmodium*, transmitted by female *Anopheles* mosquitoes [1]. Of 465 officially recognised species of *Anopheles* mosquitoes, about 70 have the capacity to transmit malaria parasites to humans [2]. In Africa, the main malaria vectors are found within four taxonomic categories: *Anopheles gambiae* complex, *Anopheles funestus* gpe, *Anopheles nili* gpe and *Anopheles moucheti* gpe [3–4]. These species complexes or groups have different distribution, behaviour and ecology [3–4].

Malaria transmission in sub-Saharan Africa is largely carried out by the *Anopheles gambiae* complex [2]. This complex comprises about nine related and morphologically indistinguishable subspecies: *Anopheles gambiae*; *Anopheles coluzzii*; *Anopheles arabiensis*; *Anopheles melas*; *Anopheles merus*; *Anopheles bwambae*; *Anopheles quadriannulatus*, *Anopheles amharicus* and *Anopheles fontaneilli* [5].

The distribution and density of these species are clearly influenced by the environmental conditions and climatic characteristics of the region [6]. *Anopheles arabiensis* is predominant in the driest areas. Although anthropophilic, it has a strong tendency to be zoophilic. However, due to the ever-changing human environment and the expansion of cities in Africa, *Anopheles arabiensis* is increasingly present in large cities [7,8]. *Anopheles melas* and *Anopheles merus* are also considered important vectors [1, 9]. *Anopheles melas* is restricted to coastal areas of West and Central Africa where its larvae develop in brackish water [10]. This species is not anthropophilic and has a short life span, making it a poor vector of malaria [11]. *Anopheles merus* is isolated in the coastal region of East Africa (1). *Anopheles gambiae* and *Anopheles coluzzii* are sympatric across much but not all of their African ranges [12]. *Anopheles coluzzii* is usually associated with permanent breeding sites, often created by human activities such as irrigation, rice cultivation and urbanisation. Whereas *Anopheles gambiae* is associated with temporary rain pools and puddles [13,14]. There is no strong evidence that the other species of the *An. gambiae* complex play a role in malaria transmission [15,16].

In the Democratic Republic of Congo (DRC), the *Anopheles gambiae* complex plays a major role in malaria transmission [17]. Most publications report a clear predominance of the former ss form, currently recognised as *Anopheles gambiae* [18–21]. *Anopheles gambiae* is widely distributed in Kinshasa [18, 20], however the Bobanga study, conducted in 2013, notes the presence of the sympatric species *Anopheles coluzzii*, in the Mont Amba district of Kinshasa [22].

Despite the high ecological diversity and the steady invasion of members of the complex into the city of Kinshasa, no updated studies on the composition and distribution of vectors of the *An. gambiae* Complex across this megacity are evident. However, the identification of species within the *An. gambiae* Complex is essential for a proper evaluation of malaria vector ecology studies and control programmes [23]. Furthermore, understanding the genetic structure of members of the *An. gambiae* Complex is important for addressing important biological and public health issues such as evolution, the spread of insecticide resistance alleles and the epidemiology of vector-borne diseases [24].

This study is proposed to establish the composition and distribution of cryptic species of the main malaria vector, *Anopheles gambiae* complex, circulating in different districts of Kinshasa city.

## Methods

### Study sites

The study was conducted in Kinshasa, during a 12-month period to include both the dry and rainy seasons (July 2021-June 2022). Kinshasa is located in Central Africa, on the right bank of the Congo River, between 4° 19′ 30’′ South latitude, and 15°19’ 20’′ East longitude. The climate is hot and humid (AW4 according to Koppen’s classification), with a rainy season from October to May and a dry season from May to September [25].

Kinshasa is a megalopolis of 9,965 km^2^. It is administratively subdivided into four districts, namely: Tshangu, Funa, Mont Amba and Lukunga. Each district is subdivided into communes. The communes are subdivided into neighbourhoods. Malaria transmission is perennial with three different strata of malaria transmission [26]. The stratum of low transmission risk, located in the north-central part of Kinshasa (malaria prevalence ≤ 5%); the stratum of intermediate transmission risk, located in the south-central part of Kinshasa (malaria prevalence between 5% and ≤ 30%) and the stratum of high transmission risk, located in the south-western and eastern part of the city of Kinshasa (malaria prevalence > 30%).

### Study design and sampling procedure

To study the distribution of members of the *An. gambiae* Complex, *Anopheles* spp. were sampled in the three malaria transmission strata within the city of Kinshasa [26]. In each stratum, two sites (or neighbourhoods) from two different communes were selected. The selection of these sites (Fig 1) was based on the administrative distribution of this megalopolis, namely four districts:

- Funa District: Kimbangu site (S 04°20’51’’S, E 15°19’12”) in the commune of Kalamu and Valée de Funa site (S04°24’49’’S, E15°16’52”), in the commune of Selembao. These two sites are located in the central-northern part of the city. It presents a variable risk of malaria transmission (prevalence ≤ 5% for the commune of Kalamu and between 5% and ≤ 30 for the commune of Selembao) [26];
- Mont Amba District: Ngwele site (S 04°20’59’’, E 15°20’17”) in the commune of Limete and Mbanza Lemba site (S 4° 24’ 59’’; E 15° 19’ 27’’) in the commune of Lemba. It presents an intermediate risk of malaria transmission (prevalence between 5% and ≤ 30%) [26];
- Lukunga District: Djelo Binza site (S 04°24’47’’, E15°20’47”) in Ngaliema commune and Kitambo site (S 04°19’37’’, E 15°16’22”) in the commune of Kitambo. These two sites are located in the western part of the city. It presents an intermediate risk of malaria transmission (prevalence between 5% and ≤ 30%) [26];
- Tshangu District, Kikimi site (S 4° 24’ 51’’; E 15° 24’ 48’’) in the commune of Kimbanseke and Ndjii Brasserie site (S 4° 28’ 35’’; E 15° 21’ 14’’): Located in the commune of Nsele. These two sites are located in the eastern part of the town. It presents a high risk of malaria transmission (prevalence > 30%) [26].

**Figure 1:**
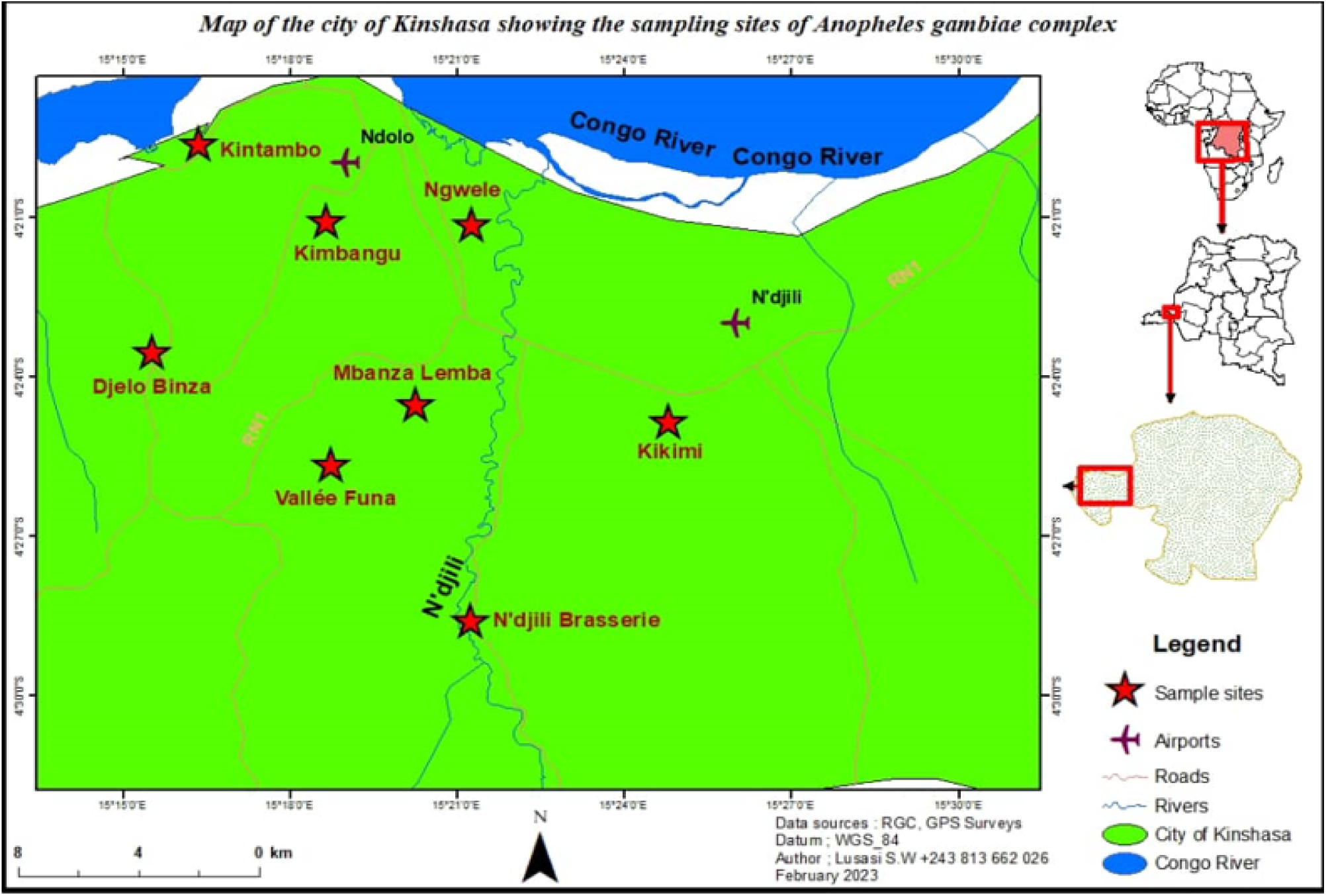
Maps of the city of Kinshasa showing the sampling sites of *Anopheles gambiae* complex.

In each study site, mosquito sampling was carried out using Center for Disease Control (CDC) light traps. In addition, typical larval sites where *Anopheles* spp. larvae develop in the different selected study sites were sampled.

### CDC Light trap catches (CDC light)

In each site, ten households were selected (by agreement of the heads of households) for collection of adult mosquitoes by CDC light trap. This collection was carried out over night between 18:00 hrs and 06:00 hrs once a month, throughout the entire study period. In each household, two traps were used, one placed inside and the other placed outside. The outdoor traps were placed within five metres of the front door. To optimise the genetic diversity of the Culicidae species collected in each study site, households selected for capture of Culicidae by CDC light trap were sampled along a 15 km long transect. The distribution of ten selected households per site followed the following pattern: three households between 0-3 km; three households between 6-9 km; and four households between 12-15 km. Different houses were used for each night, after obtaining the agreement of the heads of households. Culicidae captured were sent to the Bioecology and Vector Control Laboratory of the Kinshasa School of Public Health for morphological identification using the Coetzee key [27].

### Larva collection

The larval collection allowed us to identify the preferential larval sites of *Anopheles* spp. in the different sites of our study. In each study site, sampling was carried out twice a month during the dry season (July and August 2021) and once a month during the rainy season (between September 2021 and May 2022). An average of 200 *Anopheles* spp. larvae-positive sites were surveyed per site, along a 15 km long transect. The recognition of anopheline larvae was done by taking into account their position in relation to the water surface [27]. In order to maintain the standardisation of the data collected, only one team of two people carried out the sampling in all the sites selected, during the entire study period. All larval sites with anopheline larvae found along the determined transect were geo-referenced (Garmin GPS Map) and examined (animal hoofprints, paw prints, puddles, swamps and others) [28]. The permanent or temporary nature of the sites was assessed according to whether the larval site was connected to a water source or not.

### Molecular identification of collected samples

After morphological identification, an equal proportion of *Anopheles gambiae* complex, sampled per site, according to transect extent, were individually preserved in 1.5 mL Eppendorf tubes with silica gels and sent to the entomological research centre in Cotonou (Cotonou, Benin) for identification of cryptic species of the *An. gambiae* Complex, by polymerase chain reaction (PCR).

After extraction of the genomic DNA, the amplification and determination of the species of the *An. gambiae* complex was carried out according to the protocol of Scott [29]. For this purpose, the following primers were used:

UN: GTGTGCCGCTTCCTCGATGT

AG: CTGGTTTGGTCGGCACGTTT

AA: AAGTGTCCTTCTCCATCCTA

ME: TGACCAACCCACTCCCTTGA

QD: CAGACCAAGATGGTTAGTAT

The UN primer binds to the same position in the rDNA of all five species, and AG binds specifically to *Anopheles gambiae*. ME binds to both *Anopheles merus* and *Anopheles melas*. AA binds to *Anopheles arabiensis* and QD binds to *Anopheles quadriannulatus*.

The Santolamazza protocol allows for a better differentiation of the molecular forms of *Anopheles gambiae*, based on the specific and irreversible insertion of a 230 bp transposon (SINE200). This transposon is present on the X chromosome of *Anopheles coluzzii* whereas it is absent in *Anopheles gambiae.* This genetically inherited feature allows unambiguous, simple and direct recognition of *Anopheles gambiae* and *Anopheles coluzzii* [30]. In addition to the classical PCR components, the following primers were used:

200X6.1F-TCGCCTTAGACCTTGCGTTA and200X6.1R- CGCTTCAAGAATTCGAGATAC

### Data analysis

Data were analysed using Origin version 6.1-Scientific graphing and Data Analysis Software. Relationships between mean densities were analyzed using the t-student test. Test of significance was estimated assuming a 0.05. P-value less than 0.05 was considered significant during the analysis. Molecular diagnosis of the species was based on the primer set S200 X6.1 in *Anopheles gambiae/coluzzii*, hybrid form; Ac, *Anopheles coluzzii* (479 bp); Ag, *Anopheles gambiae* (249 bp); nc, negative control; L, scale = 100 bp (Solis BioDyne).

## Results

### Composition of *Anopheles* spp. fauna at Kinshasa and nature of identified larval sites

During the whole period of our study, 3,839 *Anopheles* were collected with the CDC light traps. After morphological identification, four species of the genus Anopheles were identified: *Anopheles gambiae* complex (11.04%), *Anopheles funestus* gpe (0.53%), *Anopheles paludis* (1.01%) and *Anopheles nili* complex (0.13%) (Fig.2).

**Figure 2:**
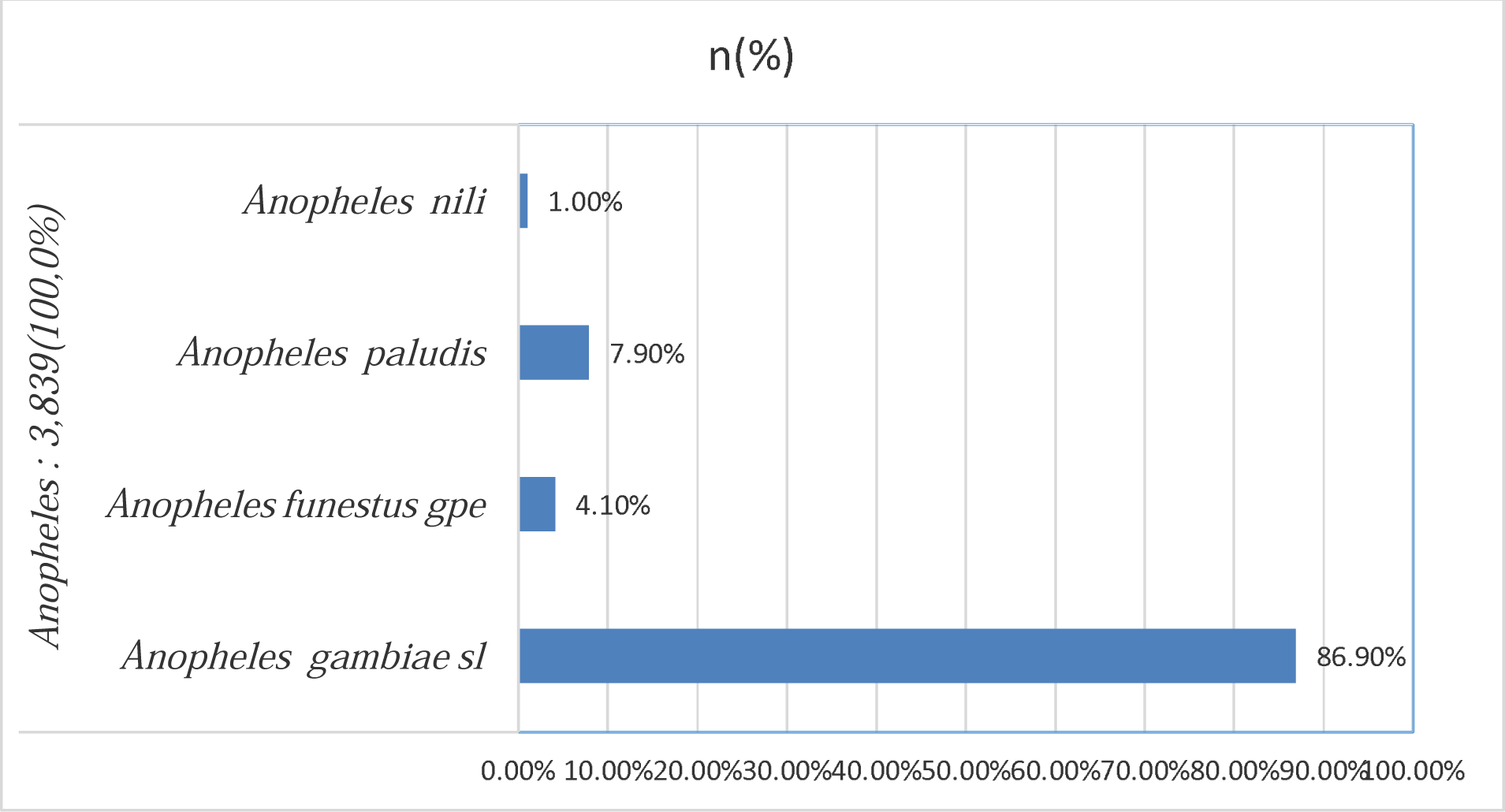
Distribution of Culicidae species from Kinshasa captured by the CDC light technique during our study.

*Anopheles gambiae* complex was identified in all sites, however, *Anopheles nili* and *Anopheles paludis* were limited in Tshangu district (Tab.1). Apart from *Anopheles paludis*, which was caught more often outside houses, the other *Anopheles* species were predominantly caught inside the sampled houses. Compared to the other *Anopheles* species identified, a significant difference was observed between the mean density of *Anopheles gambiae* complex captured by CDC light trap inside (160.33±40.41) and outside (116.50±20.17) households in Kinshasa (t = 3.36; p=0.002) (Table 2).

**Table 1.**
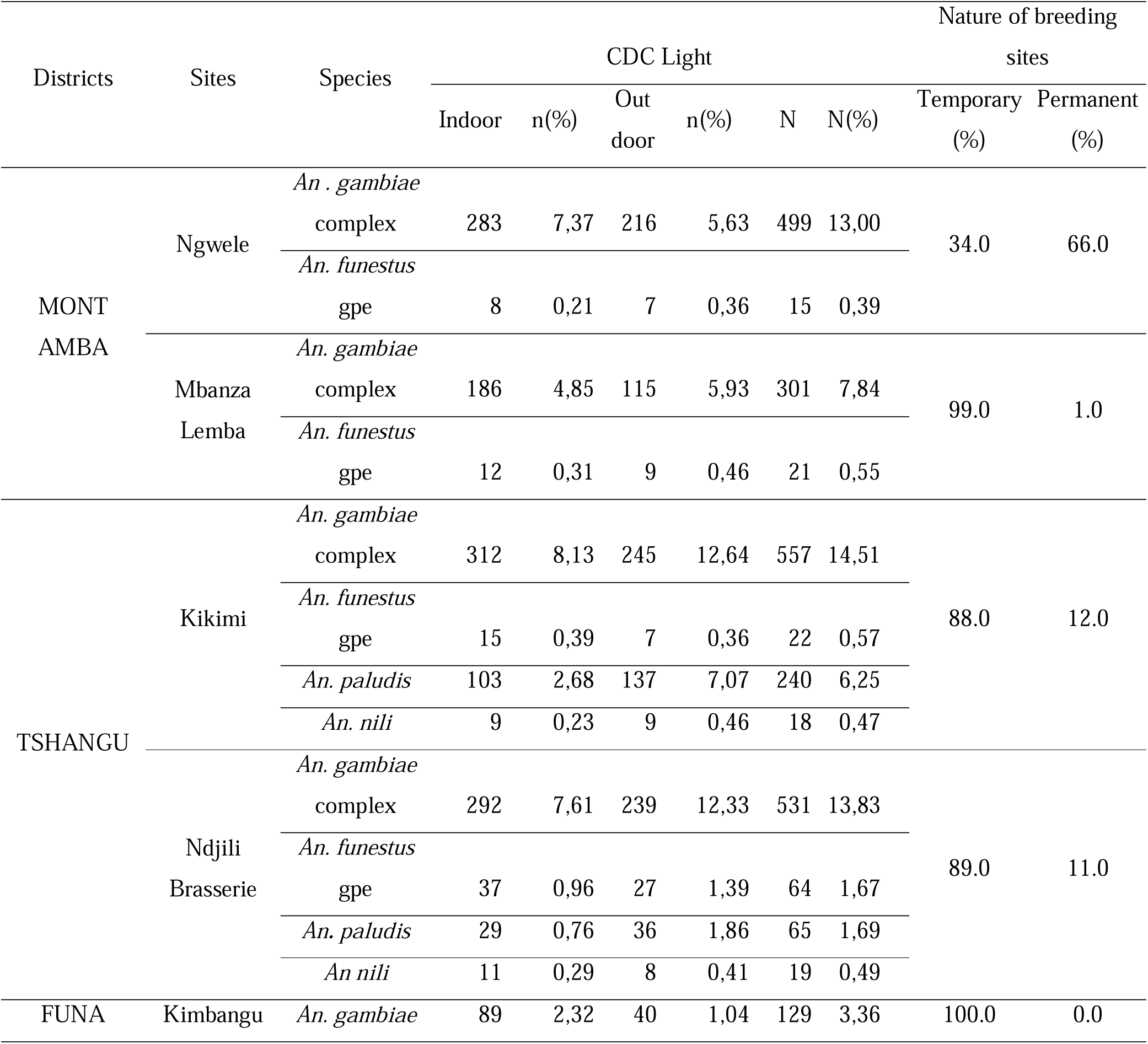
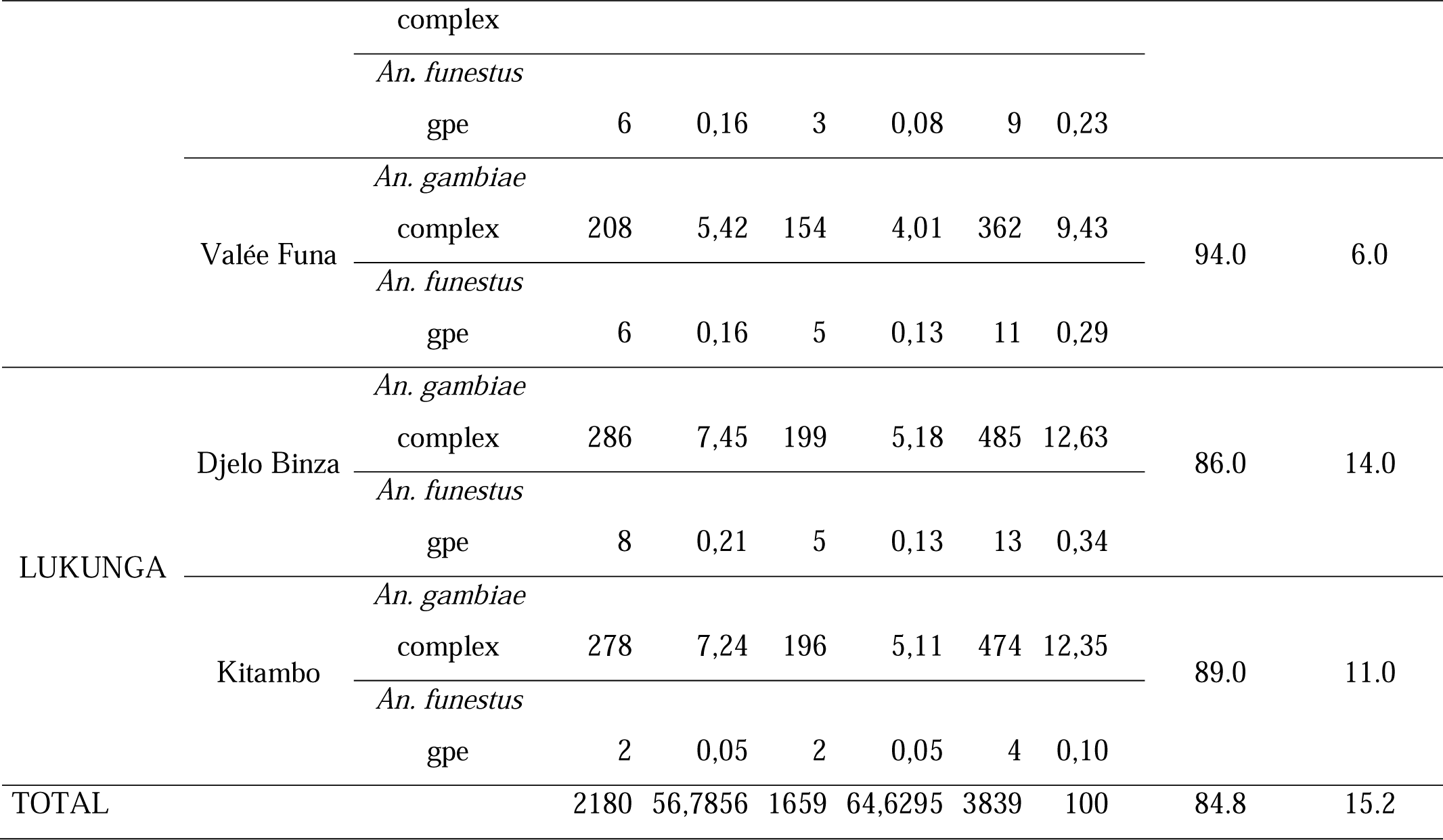
Distribution of Anopheline species captured by the CDC light technique and distribution of type of breeding sites in the four districts of Kinshasa from July 2021 to December 2022.

**Table 2.**
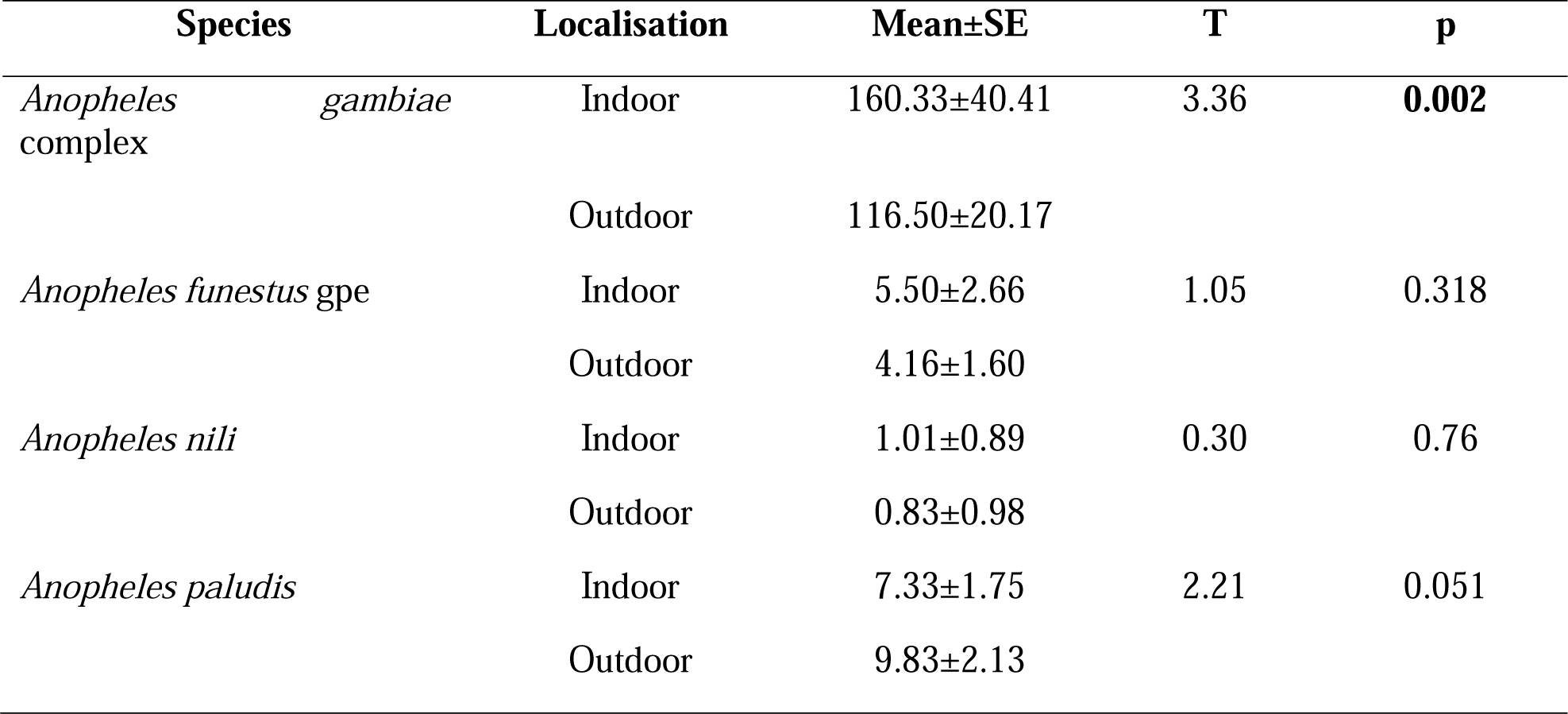
Mean indoor and outdoor density of *Anopheles* mosquitoes sampled by CDC light in Kinshasa (July-December 2021).

A diversity of larval site types was identified. Temporary sites such as puddles (water collected after rain or stagnant water from anthropogenic activities), sand extraction holes and alleys between flower beds were widely represented (81%), compared to permanent sites (rice fields, river coves, low-pressure gutters and pond edges). Permanent sites were predominant in Tshangu district compared to the other districts (Tab 1).

### Monthly dynamics of *Anopheles gambiae* complex sampled in Kinshasa

*Anopheles* spp. density dynamics were variable from month to month during the study period. The density of anopheline mosquitoes captured by CDC light was very high during the months of November and April, at 10.73% and 10.24%. Low densities were noted during July (6.69%) and February (6.20%). The monthly proportion of *Anopheles gambiae* complex caught during our study was high during November and April, 9.35% and 9.20% respectively. The peak of other *Anopheles* species, other than *Anopheles gambiae* complex, was noted in the month of January, representing 1.67% of the total *Anopheles* spp. captured (Fig 3). Results of t-student test showed that there was significant difference between the monthly means of *Anopheles gambiae* complex (280.33±60.59) and the monthly means of other *Anopheles* species (42.83±8.23) captured by CDC light during the whole study period (t=9.51; p<0.001).

**Figure 3:**
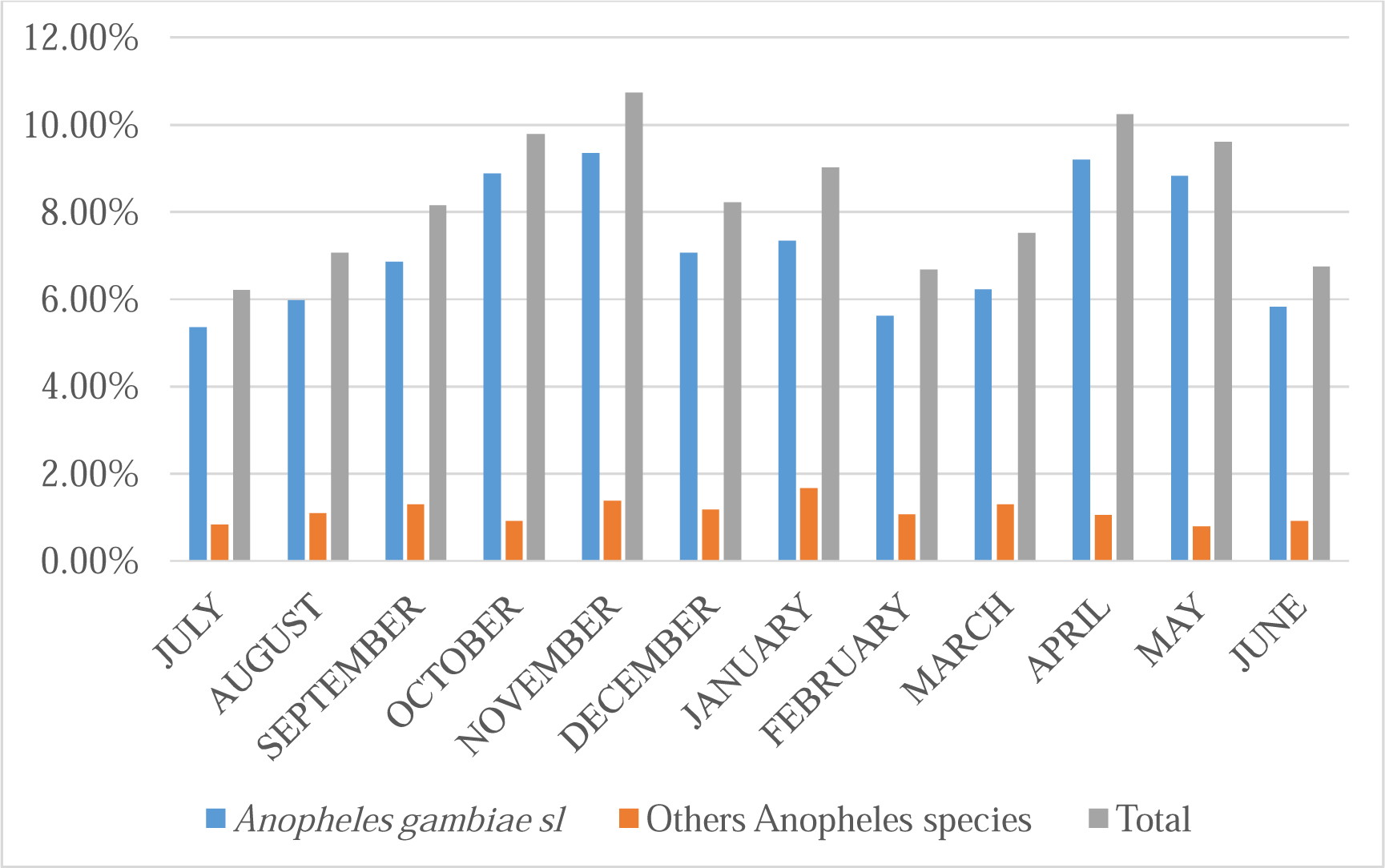
Monthly distribution of *Anopheles gambiae* complex captured by CDC light during our study.

### Molecular identification of the *Anopheles gambiae* complex collected

A total of 100 *Anopheles gambiae* complex per site, captured by CDC light, were selected. The molecular identification results of the different species of the *Anopheles gambiae* complex are presented in Table 3. Two species of this complex circulate in Kinshasa: *Anopheles gambiae* and *Anopheles coluzzii*. In all study sites, *Anopheles gambiae* was the major species, representing 674 out of 800 *Anopheles gambiae* complex that passed molecular identification.

**Table 3.**
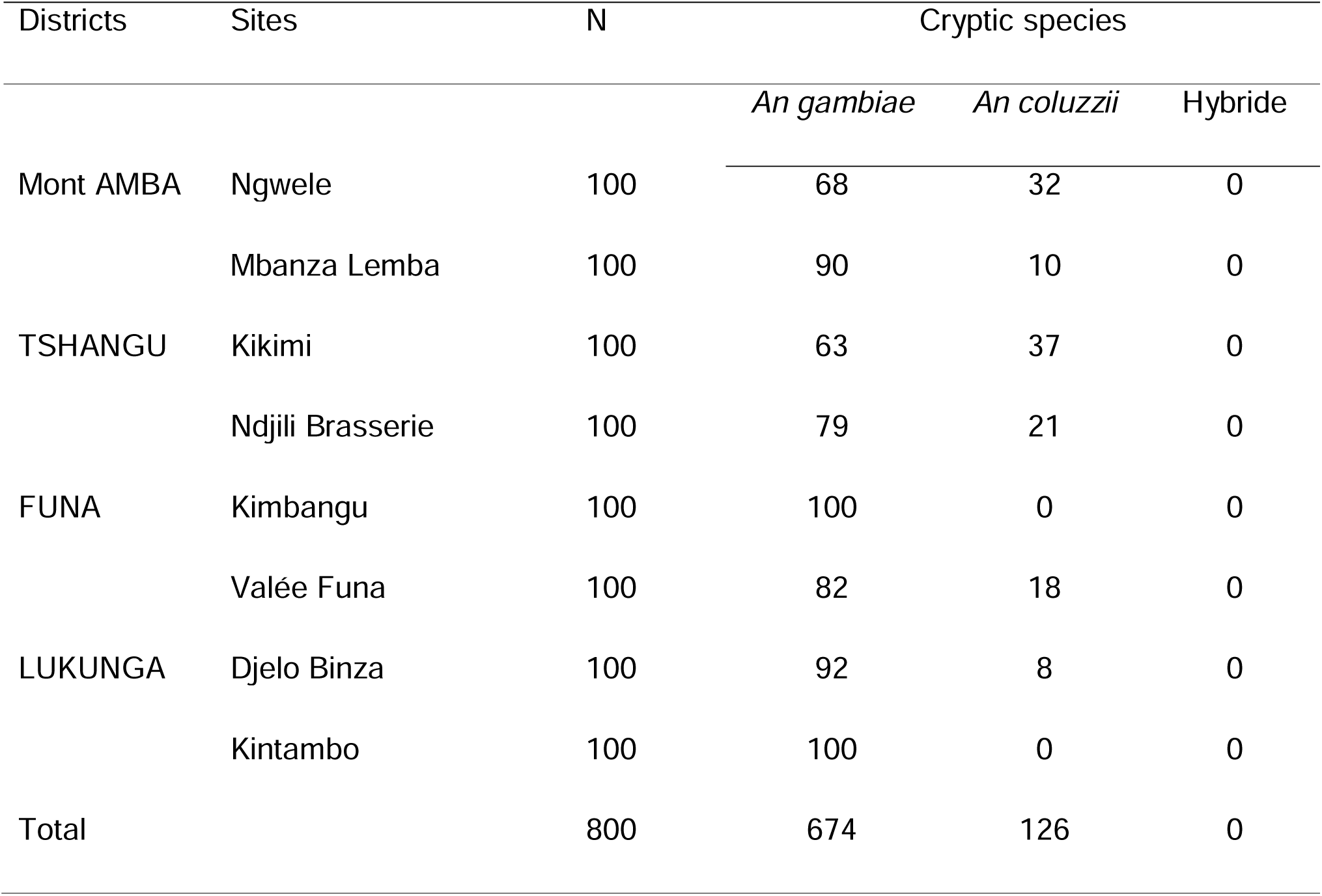
Molecular identification of cryptic species of *Anopheles gambiae* sl circulating in Kinshasa.

The sites of Kikimi and Ngwele, located in Tshangu and Mont Amba district respectively, presented a significant proportion of the total *Anopheles coluzzii* identified; 29.3 % and 25.4% respectively. *Anopheles coluzzii* was also noted in the sites of Valée de Funa (Funa District), Ndjili Brasserie (Tshangu District) and Mbanza lemba (Mont Amba District). No hybrids (*Anopheles coluzzii*/*Anopheles gambiae*) were found in the eight sites sampled for our study.

## Discussion

The identification of malaria vector species circulating in a given environment is essential for the proper evaluation of control programmes. Four species of the genus *Anopheles* were identified throughout the provincial city of Kinshasa during the period from July 2021 to June 2022. These are *Anopheles gambiae* complex, *Anopheles funestus* gpe, *Anopheles paludis* and *Anopheles nili*. Among all Anophelinae, *Anopheles gambiae* complex was the most important species, followed by *Anopheles paludis* and *Anopheles funestus* gpe. These results are consistent with the conclusions of a previous study by Karch et al [31], who found a high predominance of *Anopheles gambiae* sl among the Anopheles circulating in Kinshasa. Indeed, the high prevalence of this species of Anopheles is widely documented in urban centres on the African continent [32–33]. This can be explained by the ease with which this vector adapts to the temporary, shallow water collections that predominate in large African cities. Riveron et al, in 2018 [18]. and Zanga et al. in 2022 [20], working respectively in the east of the city of Kinshasa and across seven sites spread over the city of Kinshasa, revealed a clear predominance of *Anopheles gambiae* complex. As demonstrated by Karch [34], the *Anopheles* spp. fauna in Kinshasa is very heterogeneous and largely dominated by the presence of *Anopheles gambiae* complex. *Anopheles gambiae* is widely distributed across Africa while other species of the genus *Anopheles*, such as *Anopheles nili*, are mainly found in forest and some tropical savannah areas [35]. Indeed, *Anopheles gambiae* complex shows a preference for temporary breeding sites, which are widely distributed throughout African sub-Saharan cities and Kinshasa in particular.

In the different study sites, with the exception of *Anopheles paludis*, Anophelinae collected by CDC light trap were found in higher densities inside than outside the households. Compared to other species of the genus *Anopheles*, a significant difference was observed between the mean density of *Anopheles gambiae* complex inside and outside households (t = 3.36; p=0.002), with higher densities recorded indoors. This trend is similar to observations made by Chanyalew *et al*, in Ethiopia using the same capture technique. The higher proportion of *Anopheles gambiae* complex inside households revealed in our study can be explained by the fact that few inhabitants of Kinshasa are found outside their households after 10 pm.

Monitoring of the monthly dynamics of the genus *Anopheles* showed a fluctuation in their respective frequencies during the study period. The monthly averages were variable between species. A significant difference was noted between the monthly averages of *Anopheles gambiae* complex and the monthly averages of the other *Anopheles* species captured by CDC light during the whole study period (t=9.51; p<0.001). Compared to the other species that did not show large fluctuations, the frequencies of *Anopheles gambiae* complex increased rapidly during the rainy season, with maximum densities recorded in the months of November and April, and then decreased steadily during the dry season, with lower densities captured in the months of June and July. The fluctuation in rainfall between the dry and wet seasons in Kinshasa with the presence of temporary puddles (shallow, sunny pools) widely distributed in Kinshasa during the rainy season [20]. correlated with high larval densities of *Anopheles gambiae* complex [36–37].

Molecular identification of *Anopheles gambiae* complex captured by CDC light, showed that *Anopheles gambiae* and *Anopheles coluzzii* are the only species of the *An. gambiae* complex circulating in Kinshasa. These two species were previously identified by Riveron in and Zanga (18, 20). The approaches of these two studies were limited to very specific areas of Kinshasa. In addition, *Anopheles* sampling was based on larval collection. The choice of *Anopheles gambiae* s.l. captured by CDC light is justified by the fact that this sampling approach, targeting adult mosquitoes, best reflects the circulation of Culicidae in a given environment. The larva collection technique, although it allows a large number of samples to be obtained, is sometimes hampered by relatively low emergence rates.

*Anopheles gambiae* was in the majority in all Kinshasa districts and study sites selected. Apart from the Lukunga district, *An. coluzzii* was present throughout the city of Kinshasa. Its presence was very high in Tshangu District (Kikimi and Ndjili Brasserie sites) compared to the Mbanza lemba and Valée de Funa sites in the Mont Amba and Funa District. *Anopheles coluzzii* was also found in high proportions at the Ngwele site. Riveron *et al*, in 2018, also isolated the presence of these two sympatric species in Kinshasa, in the Tshangu district, precisely in N’Djili Brasserie.

Several studies have shown [38,39] that *Anopheles gambiae* and *Anopheles coluzzii* are the most widespread in sub-Saharan Africa. *Anopheles gambiae* and *Anopheles colu*zzii are sympatric across much of their range [12]. However, the species appear to occupy different types of breeding sites within their distribution. *Anopheles coluzzii* is generally associated with permanent breeding sites, often created by human activities such as irrigation, rice cultivation and urbanisation, whereas *Anopheles gambiae* is associated with temporary rain pools and puddles [13, 14]. Our larval sampling in Kinshasa, also found these larval site associations. *Anopheles coluzzii* was mainly present in sites where water availability favours the development of permanent breeding sites (Ngwele site with rice cultivation and Ndjili Brasserie site with a strong presence of river inlets). These associations have also been described in studies in Ghana where *Anopheles coluzzii* females were shown to preferably oviposit in permanent breeding sites [39].

Hybrid forms were not observed in the study sites selected in the four districts of Kinshasa. However, the presence of *Anopheles coluzzii* in a site with a high distribution of temporary breeding sites, such as Cité Pumbu, confirms that *Anopheles coluzzii* and *Anopheles gambiae* are distinct species but whose biological separation is far from complete [40].

Thus, studies on the ecological peculiarities of the preferred breeding sites of these two species should be very thorough in an environment undergoing considerable environmental modification and containing diverse ecological characteristics such as Kinshasa in the Democratic Republic of Congo.

### Conclusion

Understanding the epidemiology of vector-borne diseases depends on the accurate identification of known and suspected disease vectors. The present study has identified the different species of the *An. gambiae* complex present in the city of Kinshasa in the DRC, and the two species of the *Anopheles gambiae* complex present in Kinshasa, *Anopheles gambiae* and *Anopheles coluzzii,* are the main malaria vectors in Kinshasa. Two cryptic species of this complex circulate in Kinshasa. The predominance of the nature of larval sites seems to influence their distribution. Thus ongoing in depth analyses of their larval sites characteristics in Kinshasa would provide a better understanding of the influencing factors driving the distribution of members of the *An. gam*biae complex in this megalopolis.

## Data Validation and quality control

Mosquitoes were identified by experienced taxonomists using a well-established morphological key [27] and molecular techniques [29, 30]. The final dataset was validated in the Integrated Publishing Toolkit of GBIF [41]. The IPT ensures data validation through its network. Metadata fields are available at the dataset page in GBIF [42].

## Data Availability Statement

The data supporting this article are published through the Integrated Publishing Toolkit of GBIF [41] and are available under a CC0 waiver from GBIF [42].

## Re-use potential

- Previous studies on this issue have been limited to particular areas of Kinshasa. Our study has the particularity of investigating the distribution of these species throughout the city of Kinshasa.
- Secondly, it identifies the potential influence of larval habitat types on the distribution of cryptic species of Anopheles gambiae complex identified in Kinshasa.

## Competing interests

The authors declare that they have no competing interests.

## Funding

Material and financial support for this study was provided by the Bioecology and Vector Control Unit of the Environmental Health Department of the Kinshasa School of Public Health. No external funding or financial support was obtained.

## Authors’ contributions

Josue Zanga, Emery Metelo and Roger Wumba designed and implemented the study. Josue Zanga, Osee Mansiangi, Victoire Nsabatien, Maxwel Bamba and Narcisse Basosila were responsible for collecting the data. JZ, Fiacre Agossa and Nono Mvuama performed statistical analysis and prepared the manuscript for publication. All the authors helped write the manuscript. Fiacre Agossa, Degani Banzulu and Roger Wumba read and edited the manuscript before submission. The authors state that they do not have competing interests.

## Acknowledgements

We would also like to thank the technicians and researchers at the Bioecology and Vector Control Laboratory at the Kinshasa School of Public Health. We would also like to thank the molecular biology laboratory at the CREC/Benin.

## Notes

https://doi.org/10.15468/excax3

### Competing Interest Statement

The authors have declared no competing interest.

